# Comparative genome analysis revealed gene inversions, boundary expansion and contraction, and gene loss in *Stemona sessilifolia* (Miq.) Miq. chloroplast genome

**DOI:** 10.1101/2021.02.15.431246

**Authors:** Jingting Liu, Mei Jiang, Haimei Chen, Yu Liu, Chang Liu, Wuwei Wu

## Abstract

*Stemona sessilifolia* (Miq.) Miq., commonly known as Baibu, is one of the most popular herbal medicines in Asia. In Chinese Pharmacopoeia, Baibu has multiple authentic sources, and there are many homonym herbs sold as Baibu in the herbal medicine market. The existence of the counterfeits of Baibu brings challenges to its identification. To assist the accurate identification of Baibu, we sequenced and analyzed the complete chloroplast genome of *Stemona sessilifolia* using next-generation sequencing technology. The genome was 154,039 bp in length, possessing a typical quadripartite structure consisting of a pair of inverted repeats (IRs: 27,094 bp) separating by a large single copy (LSC: 81,950 bp) and a small single copy (SSC: 17,901 bp). A total of 112 unique genes were identified, including 80 protein-coding, 28 transfer RNA, and four ribosomal RNA genes. Besides, 45 tandem, 27 forward, 23 palindromic, and 72 simple sequence repeats were detected in the genome by repeat analysis. Compared with its counterfeits *(Asparagus officinalis* and *Carludovica palmate*), we found that IR expansion and SSC contraction events of *Stemona sessilifolia* resulted in two copies of *the rpl22* gene in the IR regions and partial duplication of the *ndhF* gene in the SSC region. Secondly, an approximately 3-kb-long inversion was identified in the LSC region, leading to *the petA* and *cemA* gene presented in the complementary strand of the chloroplast DNA molecule. Comparative analysis revealed some highly variable regions, including *trnF-GAA_ndhJ, atpB_rbcL, rps15_ycf1, trnG-UCC_trnR-UCU, ndhF_rpl32.* Finally, gene loss events were investigated in the context of phylogenetic relationships. In summary, the complete plastome of *Stemona sessilifolia* will provide valuable information for the molecular identification of Baibu and assist in elucidating the evolution of *Stemona sessilifolia.*

## Introduction

Radix Stemonae, also known as Baibu, is one of the most popular herbal medicines used in many Asian countries, including China, Korea, Japan, Thailand, and Vietnam. It has been used in treating various respiratory diseases such as bronchitis, pertussis, and tuberculosis [1,2]. It was also well known for killing cattle parasites, agricultural pests, and domestic insects [3, 4]. Stenine B, one of the major chemical ingredients of Baibu, has been considered a potential drug candidate against Alzheimer’s disease due to its significant acetylcholinesterase inhibitory activity [5]. Owing to the important medicinal values, extensive genetic, biochemical, and pharmacological studies on Baibu is needed.

According to Pharmacopoeia of the People’s Republic of China (2015 edition), the root tubers of *Stemona tuberosa*, *Stemona japonica*, and *Stemona sessilifolia* were all considered as the authentic sources of Baibu. Although these three species were all employed as the raw materials of Baibu, we cannot ignore their inherent difference. For example, Stemona alkaloids are the major components responsible for Baibu’s antitussive activities. However, their composition and contents vary among *S. tuberosa, S. japonica,* and *S. sessilifolia* [6, 7]. These three species differ in antitussive, anti-bacterial, and insecticidal activities [8]. Therefore, it is critical to determine the exact origin of plant materials used as Baibu.

On the other hand, multiple authentic sources and the homonym also increase the difficulty of identifying Baibu. In some area of China, another herbal medicine, *Aconitum kusnezoffii* Rchb., is also called Baibu. However, the therapeutic activity of *Aconitum kusnezoffii* is significantly different from the authentic sources of Baibu described in Chinese Pharmacopoeia. Researches even reported that it might result in toxicity when *Aconitum kusnezoffii* was taken in large quantities [9]. Besides, counterfeits in the herbal market also brought challenges to the exact identification of Baibu. Due to their similar morphologic features to the authentic sources for Baibu, many counterfeits such as *Asparagus officinalis, Asparagus filicinus,* and *Asparagus acicularis* were sold as Baibu in the herbal market frequently [10]. Therefore, the exact identification of Baibu origin is critical for its usage as a medicinal herb.

DNA barcode was deemed a more efficient and effective method in identifying plant species compared to morphological characteristics. Typical barcodes such as *ITS, psbA-trnH, matK,* and *rbcL* have been used to distinguish different plant species [11–13]. However, these DNA barcodes were not always working effectively, especially when distinguishing closely related plant species. Such a phenomenon may attribute to single-locus DNA barcodes still lack adequate variations in closely related taxa. Compared with DNA barcodes, the chloroplast genome provides more abundant genetic information and higher resolution in identifying plant species. Some researchers have proposed using the chloroplast genome as a species-level DNA barcode [14, 15].

The chloroplast is an organelle presenting in almost all green plants. It is the center of photosynthesis and plays a vital role in sustaining life on earth by converting solar energy to carbohydrates. Besides photosynthesis, chloroplast also plays critical roles in other biological processes, including the synthesis of amino acids, nucleotides, fatty acids, and many secondary metabolites. Furthermore, metabolites synthesized in chloroplasts are often involved in plants’ interactions with their environment, such as response to environmental stress and defense against invading pathogens [16–18]. Due to its essential roles in the cellular processes and relatively small genome size, the chloroplast genome is a good starting point for resolving phylogenetic ambiguity, discriminating closely related species, and revealing the plants’ evolutionary process. To date, over 5000 chloroplast genomes from a variety of land plants are available. Phylogenetic analyses have demonstrated chloroplast genomes’ effectiveness in inferring phylogenetic and distinguishing closely related plant species [19, 20].

Unfortunately, the taxonomic coverage of the sequenced chloroplast genome is somewhat biased. For example, until now, the chloroplast genome of *Stemona sessilifolia* has not been reported. The lack of chloroplast genome information prohibited studies aiming to understand the evolutionary processes in the family Stemonaceae. Here, we reported the full plastid genome of *Stemona sessilifolia.* Based on the sequence data, we performed a multi-scale comparative genome analysis among *Stemona sessilifolia, Asparagus officinalis,* and *Carludovica palmate* (the major counterfeits of Baibu). We investigated the difference among these three species from three aspects, including general characteristics, repeat sequences, and sequence divergences. We also characterized the significant changes, including genome rearrangement, IR expansion, and SSC contraction, in the plastid genome of *Stemona sessilifolia, Asparagus officinalis,* and *Carludovica palmate*.

Lastly, we investigated the gene loss events in Stemonaceae and its closely related families (Asparagoideae, Velloziaceae, Cyclanthaceae, Pandanaceae). The results obtained in this work will provide valuable information for species identification of herb materials that are used as Baibu. Furthermore, it lays the foundation for elucidating the evolutionary history of plant species in the family Stemonaceae.

## Materials and Methods

### Plant Material and DNA Extraction

We collected fresh young leaves of *Stemona sessilifolia* from multiple individuals in the Institute of Medicinal Plant Development (IMPLAD), Beijing, China, and stored them at −-80°C for chloroplast DNA extraction. All samples were identified by Professor Zhao Zhang, from the Institute of Medicinal Plant Development, Chinese Academy of Medical Sciences & Peking Union Medical College. The voucher specimens were deposited in the herbarium of IMPLAD. *Stemona sessilifolia* is not an endangered or protected species. Therefore, specific permissions for the collection of *Stemona sessilifolia* were not required. Total DNA was acquired from 100mg fresh young leaves using a plant genomic DNA kit (Tiangen Biotech, Beijing, Co., Ltd.). Finally, 1.0% agarose gel and Nanodrop spectrophotometer 2000 (Thermo Fisher Scientific, United States) was used to evaluate the purity and concentration, respectively.

### Genome Sequencing, Assembly, and Annotation

According to the standard protocol, the DNA of *Stemona sessilifolia* was sequenced using the Illumina Hiseq2000 platform, with insert sizes of 500 bases for the library. A total of 5,660,432 paired-end reads (2 × 250bp) were obtained, and low-quality reads were trimmed with Trimmomatic software [21].

To extract reads belonging to the chloroplast genome, we downloaded 1,688 chloroplast genome sequences from GenBank and constructed a Basic Local Alignment Search Tool (BLASTn) database. All trimmed reads were mapped to this database using the BLASTN program [22], and reads with E-value > 1E-5 were extracted. The reads were assembled first using the SPAdes software with default parameters [23]. The contigs were then subjected to gap closure using the Seqman module of DNASTAR (V11.0) [24]. Finally, we evaluated the assembled genome’*s quality* by mapping the reads to the genome using Bowtie2 (v2.0.1) with default settings [25]. For further evaluation, all the barcode sequences of *Stemona sessilifolia* available in GeneBank were download (S1 file), including *matK* (1), *petD(1), rbcL* (1), *rpoC1* (1), *rps16* (1), *rps19-rpl22-psbA* (1), *trnL* (3), and *trnL-trnF* (2), the number enclosed in parentheses represented the number of barcode sequence. The BLAST program was used to calculate the identity between the chloroplast genome sequence of *Stemona sessilifolia* and each barcode sequence. As a result, the barcode of rps19-rpl22-psbA is located at the boundary of LSC/IRb, with an identity value of 100%. All the other barcode sequences also gave identity values of 100%, indicating the high reliability of the chloroplast genome sequence.

Gene annotation of *Stemona sessilifolia* chloroplast genome was conducted using the CpGAVAS web service with the default parameters [26]. The tRNA genes were confirmed with tRNAscan-SE [27] and ARAGORN [28]. Then the gene/intron boundaries were inspected and corrected using the Apollo program [29]. The Cusp and Compseq programs from EMBOSS were used to calculate the codon usage and GC content [30]. Finally, OrganellarGenomeDRAW [31] was used to generate the circular chloroplast genome map of *Stemona sessilifolia.*

### Repeat Sequence Analysis

Perl script MISA(http://pgrc.ipk-gatersleben.de/misa/) was used to identify simple sequence repeats (SSRs) with the following parameters: 8 repeat units for mononucleotide SSRs, 4 repeat units for di- and tri-nucleotide repeat SSRs, and 3 repeat units for tetra-, Penta-, and hexanucleotide repeat SSRs. Tandem Repeats Finder was used with parameters of 2 for matches and 7 for mismatches and indels [32]. For the minimum alignment score and the maximum period, the size was set to 50 and 500. Palindrome and forward repeats were identified by the REPuter web service [33]. The minimum repeat size and the similarity cutoff were set to 30 bp and 90%, respectively.

### Comparative Genomic Analysis

A total of four species, including *Stemona sessilifolia, Asparagus officinalis* (NC_034777), *Carludovica palmate* (NC_026786), and *Sciaphila densiflora* (NC_027659), were subjected to multiple sequence alignment using mVISTA with default parameters [34]. Subsequently, 20 introns and 108 intergenic regions shared by *Stemona sessilifolia, Asparagus officinalis,* and *Carludovica palmates* were extracted using custom MatLab scripts to perform sequence divergence analysis. Firstly, sequences of each intergenic-region/intron were aligned individually using the CLUSTALW2 (v2.0.12) [35] program with options “-type = DNA-gapopen = 10-gapext = 2”. Secondly, Pairwise distances were calculated with the Distmat program in EMBOSS (v6.3.1) using the Kimura 2-parameters (K2p) evolution model [36]. We attempted to discover highly divergent regions for the development of novel molecular markers. To identify the occurrence of genome rearrangement events in the chloroplast genome of *Stemona sessilifolia,* synteny analysis among the three species mentioned above were performed using Mauve Alignment [37].

### Phylogenetic Analysis

A total of 11 chloroplast genomes were distributed into Stemonaceae (3), Cyclanthaceae (1), Pandanaceae (1), Velloziaceae (1), and Asparagoideae (5) were retrieved from the RefSeq database. The protein sequences shared by these chloroplast genomes were used to construct a phylogenetic tree with *Veratrum patulum* and *Paris dunniana* as outgroup taxa (S1 Table). Fifty-eight proteins were involved, including ACCD, ATPA, ATPB, ATPE, ATPF, ATPH, ATPI, CLPP, MATK, NDHB, NDHC, NDHJ, NDHK, PETA, PETB, PETD, PETG, PETL, PETN, PSAA, PSAB, PSAJ, PSBA, PSBB, PSBC, PSBD, PSBE, PSBF, PSBH, PSBI, PSBJ, PSBK, PSBL, PSBM, PSBN, PSBT, RBCL, RPL2, RPL14, RPL16, RPL22, RPL23, RPL33, RPL36, RPOA, RPOB, RPOC1, RPS2, RPS3, RPS4, RPS7, RPS8, RPS11, RPS14, RPS18, RPS19, YCF3, AND YCF4. All these protein sequences were aligned using the CLUSTALW2 (v2.0.12) program with options “-gap open = 10-gapext = 2-output = phylip”. We used Maximum Likelihood (ML) method to infer the evolutionary history of *Stemona sessilifolia* and species closely related to it. The detailed parameters were “raxmlHPC-PTHREADS-SSE3 -f a -N 1000 -m PROTGAMMACPREV –x 551314260 -p 551314260 -o Nicotiana_tabacum, Solanum_lycopersicum -T 20”.

## Results and discussion

### General characteristics of chloroplast genomes

The gene map of *Stemona sessilifolia* is shown in Fig 1. The sequence is provided in S2 File along with those of the major counterfeit of Baibu, *Asparagus officinalis* (NC_034777), and *Carludovica palmate* (NC_026786). The chloroplast genomes of *Stemona sessilifolia* and two other species share the standard features of possessing a typical quadripartite structure consisting of a pair of inverted repeats (IRs) separating a large single copy (LSC) and a small single copy (SSC), similar to other angiosperm chloroplast genomes [38].

**Figure 1.**
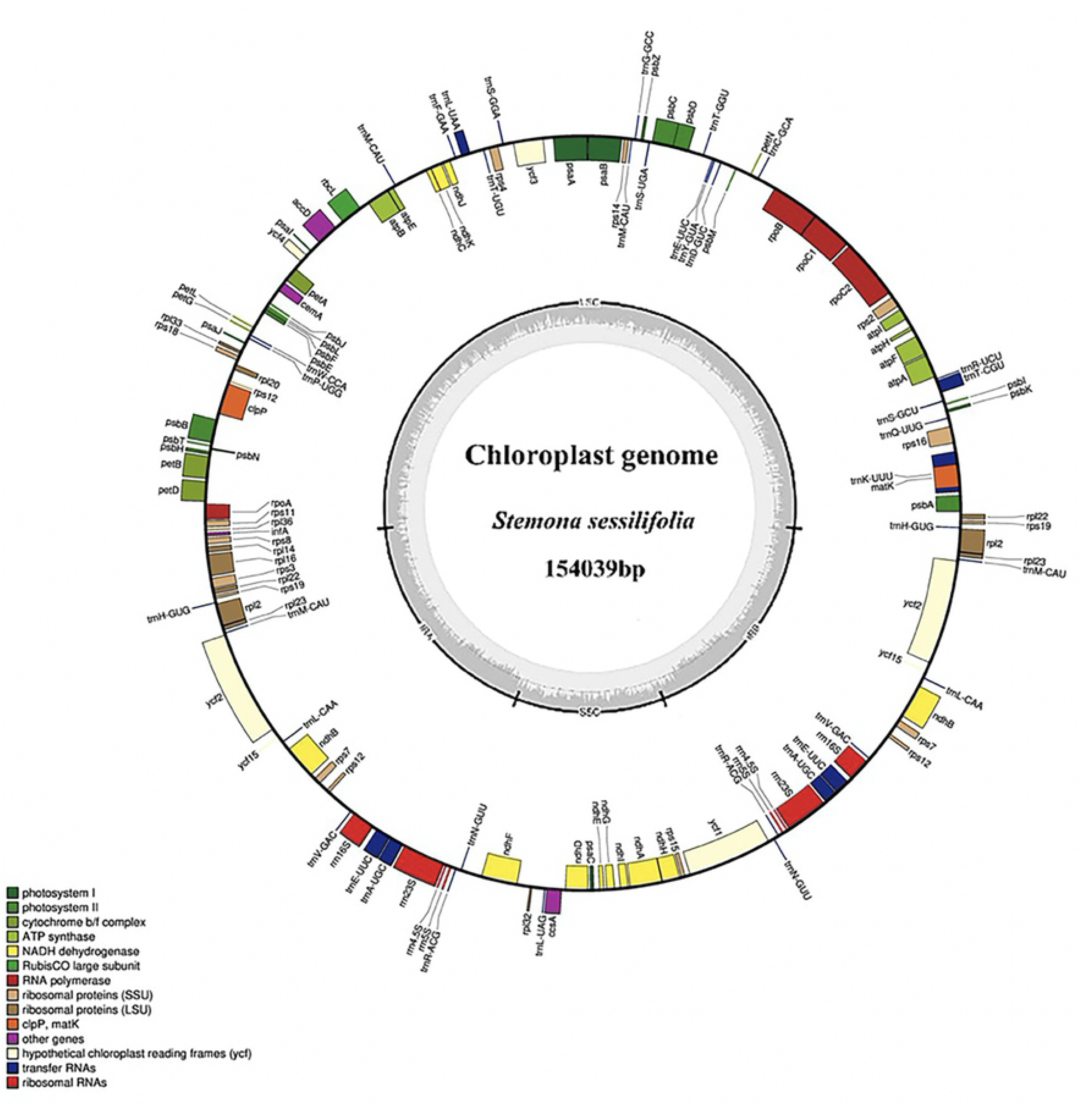
Gene maps of chloroplast genomes of *Stemona sessilifolia, Asparagus officinalis, and Carludovica palmate.* Genes inside and outside the circle were transcribed clockwise and counterclockwise, respectively. The darker gray in the inner circle indicated GC content. Genes with different functions were characterized with varying bars of color

We then carried out a multi-scale comparative genome analysis of these three chloroplast genomes from four aspects, including the size, the guanine-cytosine (GC) content, the count of genes, and the gene organization (Table 1). The complete circle chloroplast genomes of *S. sessilifolia, A. officinalis,* and *C. palmate* were 154,039 bp, 156,699 bp, and 158545 bp, respectively. Compared to *A. officinalis* and *C. palmate, S. sessilifolia* showed a relatively short SSC region and a relatively long IR region. We speculated that the chloroplast genome of *S. sessilifolia* might undertake IR expansion and SSC contraction simultaneously. There has no significant difference among *S. sessilifolia, A. officinalis,* and *C. palmate.* Such a result may attribute to the high conservation of tRNAs and rRNAs. The length of CDS regions of *A. officinalis* and *C. palmate* is shorter than *S. sessilifolia,* indicating gene loss events may occur in the chloroplast genome of *A. officinalis* and *C. palmate.*

**Table 1.** Chloroplast genome characteristics of *Stemona sessilifolia Asparagus officinalis* and *Carludovica palmate.*

| Plastome | Characteristics | Species |  |  | State |
| --- | --- | --- | --- | --- | --- |
|  |  | *Stemona sessilifolia* | *Asparagus officinalis* | *Carludovica palmate* |  |
| Size (bp) | Genome | 154039 | 156699 | 158545 | > |
|  | LSC | 81950 | 84999 | 71426 | > |
|  | IR | 27094 | 26531 | 26529 | < |
|  | SSC | 17901 | 18638 | 18364 | > |
|  | tRNA genes | 2874 | 2863 | 2816 | < |
|  | rRNA genes | 9056 | 9052 | 8866 | < |
|  | CDS | 79641 | 77436 | 77802 | < |
| GC content | Overall | 38.00 | 37.59 | 37.74 | < |
|  | LSC | 36.18 | 35.60 | 35.79 | < |
| (%) | IR | 42.70 | 42.92 | 42.81 | > |
|  | SSC | 32.13 | 31.50 | 31.51 | < |
|  | tRNA genes | 53.42 | 53.57 | 53.40 | > |
|  | rRNA genes | 55.22 | 55.38 | 55.38 | - |
|  | CDS | 38.31 | 38.1 | 38.41 | > |
|  | 1st position | 45.7 | 45.64 | 45.93 | > |
|  | 2nd position | 38.46 | 38.56 | 38.39 | > |
|  | 3rd position | 30.78 | 30.09 | 30.91 | < |
| NO. of genes | Total | 112 | 110 | 112 | < |
|  | protein-coding genes | 80 | 78 | 80 | < |
|  | tRNAs | 28 | 28 | 28 | - |
|  | rRNAs | 4 | 4 | 4 | - |
|  | Genes with introns | 18 | 18 | 18 | - |
|  | Genes in IR | 21 | 22 | 18 | > |

LSC: Large single-copy, IR: Inverted repeat, SSC: Small single-copy, CDS: Coding sequence. ">" and "<" indicated the characteristic parameters of *Asparagus officinalis* greater than and less than *Stemona sessilifolia*, respectively. "-" represented the characteristic parameters of *Asparagus officinalis* and *Stemona sessilifolia* were equal to each other.

For GC content, *S. sessilifolia* showed a higher value in LSC, SSC, and CDS regions than *A. officinalis* and *C. palmate,* even in the complete chloroplast genome. However, in the IR regions, *A. officinalis* and *C. palmate* showed a GC content value larger than *S. sessilifolia.* The GC content decreased remarkably from the first position to the third position in the codon position scale. Such a result was in line with the phenomenon observed in most land plant plastomes.

We identified 112, 110, and 112 genes in the chloroplast genomes of *S. sessilifolia, A. officinalis,* and *C. palmate,* respectively. All of these three chloroplast genomes have 28 tRNAs and four rRNAs. The number of genes with introns in each species is 18, similar to reports in prior works [39]. Therefore, we may conclude that there have no intron loss events occurred in the chloroplast genomes of these three species. All the genes with introns were described in Table S2. Besides, 21, 22, and 18 genes were predicted for *S. sessilifolia, A. officinalis,* and *C. palmate* in IR regions.

The gene organization*s* were compared in Table 2. In the upstream region and the downstream region of the *C. palmate* chloroplast genome, premature stop codons were discovered in the *ycf1* gene, resulting in the loss of this gene. Compared to *S. sessilifolia,* we found the shorter CDS regions of *C. palmate* is directly related to the loss of this gene. We also found a full-length and a pseudogene of *ndhF* gene coexist in the chloroplast genome of *S. sessilifolia,* which further indicated SSC contraction events.

**Table 2.**
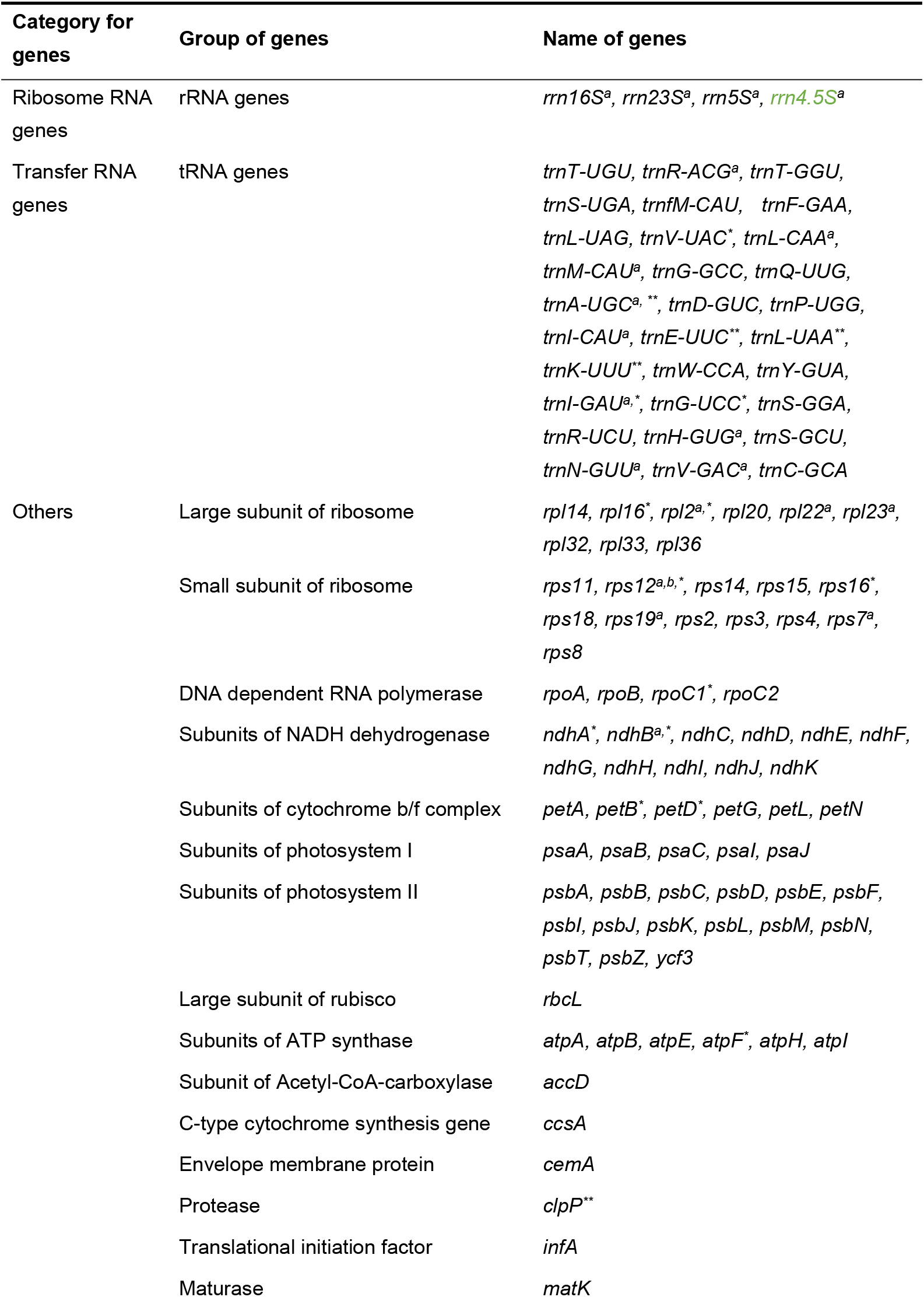
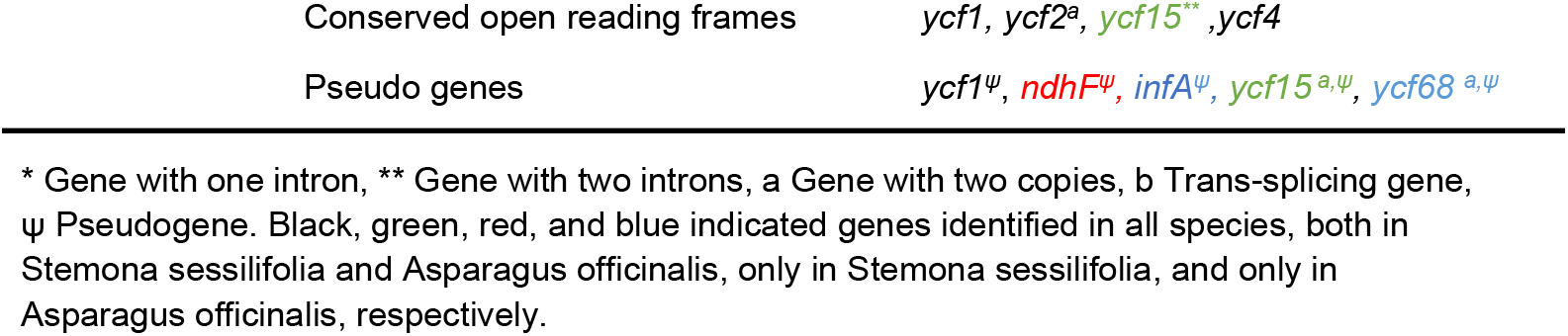
Genes presented in chloroplast genomes of *Stemona sessilifolia, Asparagus officinalis* and *Carludovica palmate.*

### Repeat Sequence Analysis

Simple sequence repeats (SSRs), the tandem repeat sequences consisting of 1-6 repeat nucleotide units, are widely distributed in prokaryotic and eukaryotic genomes. High polymorphism makes the SSRs effective molecular markers in species identification, population genetics, and phylogenetic research [40, 41]. In the current study, we investigated the distribution of SSRs in the genomes and their count and type (Fig 2). As a result, a total of 81,59, and 72 SSRs were detected in *S. sessilifolia, A. officinalis,* and *C. palmate,* respectively. Mononucleotide motifs showed the highest frequency of SSRs in these species, followed by di-nucleotides and tri-nucleotides. Compared to *A. officinalis* and *C. palmate, S. sessilifolia* contained more SSRs. However, three tri-nucleotide repeats were detected in *C. palmate* and one in *A. officinalis,* but none were identified in *S. sessilifolia.* As expected, most mono-nucleotide and di-nucleotide repeats consisted of A/T and AT/AT repeats, respectively. The results suggest that these chloroplast genomes are rich in short poly-A and poly-T motifs, while poly-C and poly-G are relatively rare. We then use Tandem Repeats Finder [32] and REPuter [33] to detect long repeats and found 95, 70, and 95 long repeat sequences in *S. sessilifolia*, *A. officinalis*, and *C. palmata,* respectively. For *S. sessilifolia*, the number of Tandem repeats, Forward repeats, and Palindromic repeats was 45, 27, and 23, respectively. The number of the corresponding repeat sequences for *A. officinalis* was 45, 11, and 14, respectively. The number of the repeat sequences for *C. palmata* was 45, 33, and 17, respectively.

**Figure 2.**
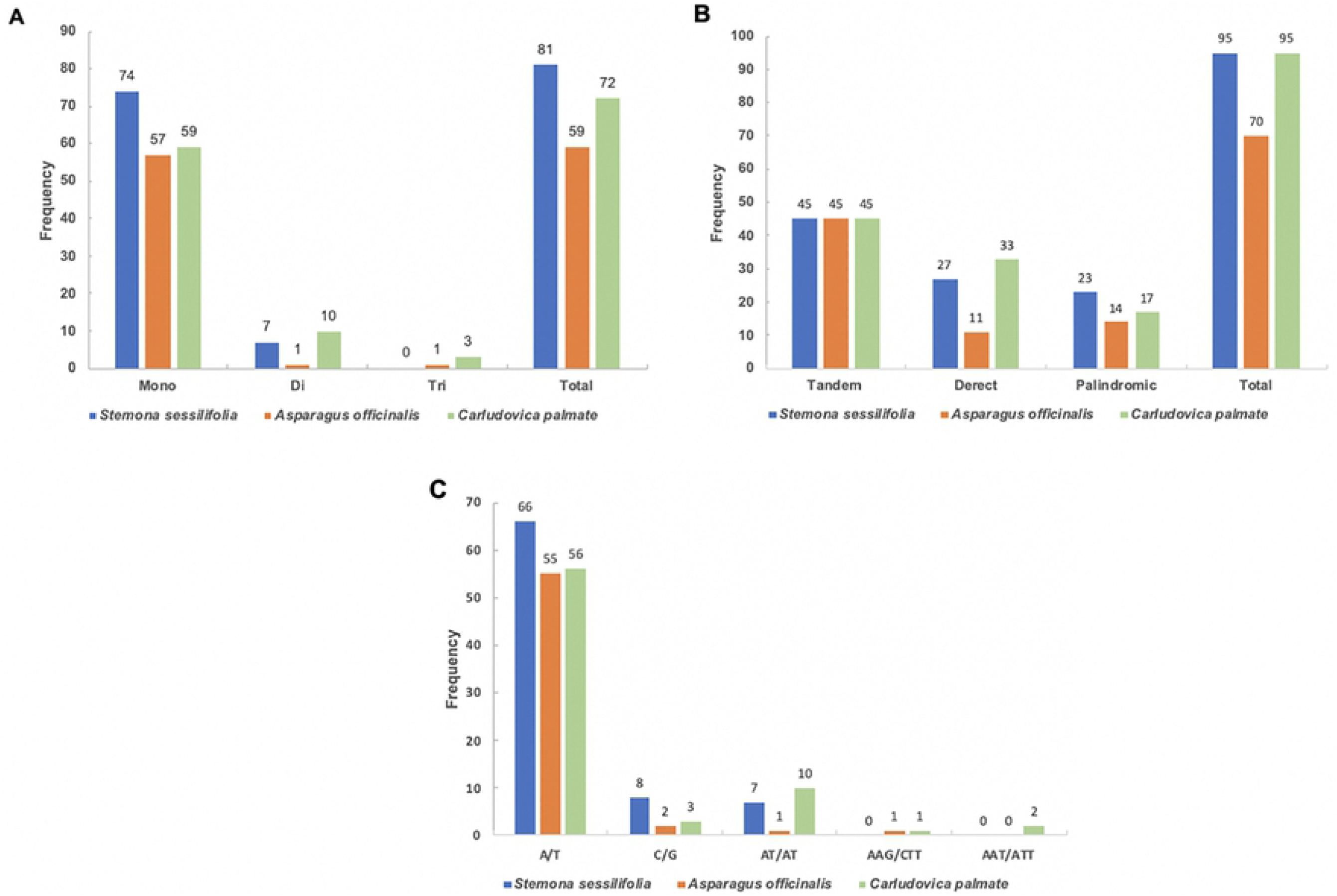
Simple sequence repeats (SSRs) and long repeat sequences are identified in the chloroplast genomes. (A) Distribution of different types of SSRs in the chloroplast genomes. (B) Distribution of long repeat sequences in the chloroplast genomes. (C) Frequency of SSR motifs in different repeat class types.

There have significant differences in the types of repeat sequence among *S. sessilifolia, A. officinalis,* and *C. palmata.* The repeat occurrence in *S. sessilifolia was* similar to that of *A. officinalis* but significantly higher than that of *C. palmata.* It should be noted that the size of *the A. officinalis* and *C. palmata* chloroplast genome is larger than the chloroplast genomes of *S. sessilifolia.* Therefore, the relatively larger size of the chloroplast genome of *A. officinalis* and *C*. *palmata* does not result from the repeat sequence.

### Sequence divergence analysis

To evaluate the genome sequence divergence, we aligned sequences from four species using mVISTA [34] (Fig 3). The chloroplast genome of *S. sessilifolia* was significantly different from *A. officinalis* and *C. palmata.* Severe gene loss events always lead to highly reduced plastomes [20, 42]. As expected, the non-coding regions were more divergent than coding regions among these species. The two most divergent regions were *ycf4-psbJ* regions (red square A) and *rpl22* coding regions (red square B). We suspected that such a phenomenon might result from gene loss events or genome rearrangement events, and the detailed reasons will be discussed later. *Ycf1* gene is also highly divergent, which may occur due to the occurrence of pseudogenization. In summary, the LSC region showed the highest divergence, followed by the SSC region, and the IR regions were less divergent than the LSC and SSC region. Compared to the coding areas, the intergenic spacers displayed higher divergence.

**Figure 3.**
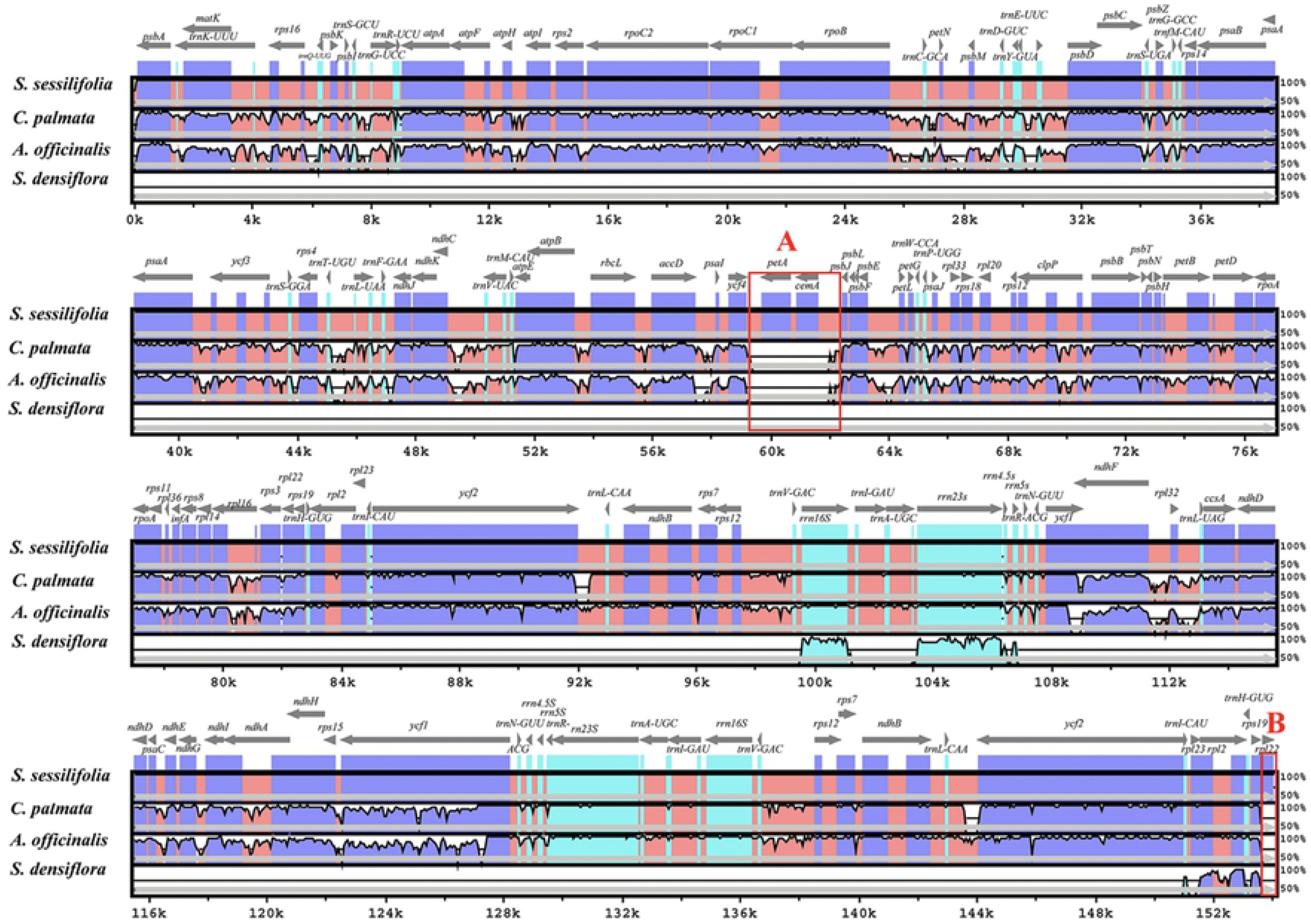
Comparison of four chloroplast genomes using mVISTA program. Gray arrows indicated the orientations and positions of genes. Untranslated regions, conserved non-coding regions, and coding regions were characterized by sky-blue block, red block, and blue block. We adopted a cutoff value of 70% in the process of alignment.

Highly divergent regions always assist in the development of molecular markers. Because non-coding regions are evolved more rapidly than coding regions, the intergenic regions and intron regions were always considered ideal candidate regions of molecular markers with high resolution. Therefore, we calculated the Kimura 2-parameter (K2p) distances for each set of the intergenic regions and intron regions. A relatively higher K2p value between any two species is necessary to distinguish each species from the other two species. Therefore, we calculated the minimal K2P (MK2P) value for each set of intergenic regions and intron regions. The non-coding regions with higher MK2P values are likely to be the candidate regions of high-resolution molecular markers. Consequently, for introns (S3 Table), the MK2p value ranges from 0.0055 to 0.1096. *ClpP*_intron2 with the highest MK2p value followed by *rpl16*_intron1, the third, fourth, and fifth were *rps16*_intron1, *ndhA*_intron1, and *trnL-UAA_intron1,* respectively. For intergenic spacers (S4 Table), five highly conserved intergenic spacers were observed, including *ndhA_ndhH, psaB_psaA, psbL_psbF, rpl2_rpl23,* and *trnI-GAU_trnA-UGC.* The MK2p value of intergenic spacers ranges from 0 to 0.3301, and the top-10 intergenic spacers with higher MK2p values were listed as follows: *trnF-GAA_ndhJ, atpB_rbcL, rps15_ycf1, trnG-UCC_trnR-UCU, ndhF_rpl32, accD_psaI, rps2_rpoC2, trnS-GCU_trnG-UCC, trnT-UGU_trnL-UAA,* and *rps16_trnQ-UUG.* In conclusion, compared to introns, we observed higher divergence in intergenic spacers. The intergenic spacers with large K2p values represent good candidate molecular markers to distinguish these three species.

### Rearrangement of the chloroplast genome

To investigate whether there are significant differences in *ycf3-psbJ* regions (red square A in Fig 3) and *rpl22* coding regions (red square B in Fig 3) between *S. sessilifolia* and its closely related species, we conducted synteny analysis. As plotted in Fig 4, we detected a large inversion of 3 kb long in the LSC region. Interestingly, such an approximately 3-kb long inversion was confirmed located in *ycf3-psbJ* regions. Therefore, we can conclude that the occurrence of genome rearrangement events leads to a significant difference in *the ycf3-psbj* areas between *S. sessilifolia* and the other two species. To investigate whether such an inversion that exists in *S. sessilifolia* is unique, we conducted synteny analysis between the chloroplast genome of *S. sessilifolia* and species in Lioscoreales and Liliales, which belong to the two closely related orders of Pandanales. Compared to any species in Dioscoreales and Liliales, inversion in *ycf3-psbJ* regions in *S. sessilifolia* was always visible (data are not shown). Therefore, inversion in the *ycf3-psbj* areas may be unique to *S. sessilifolia*.

**Figure 4.**
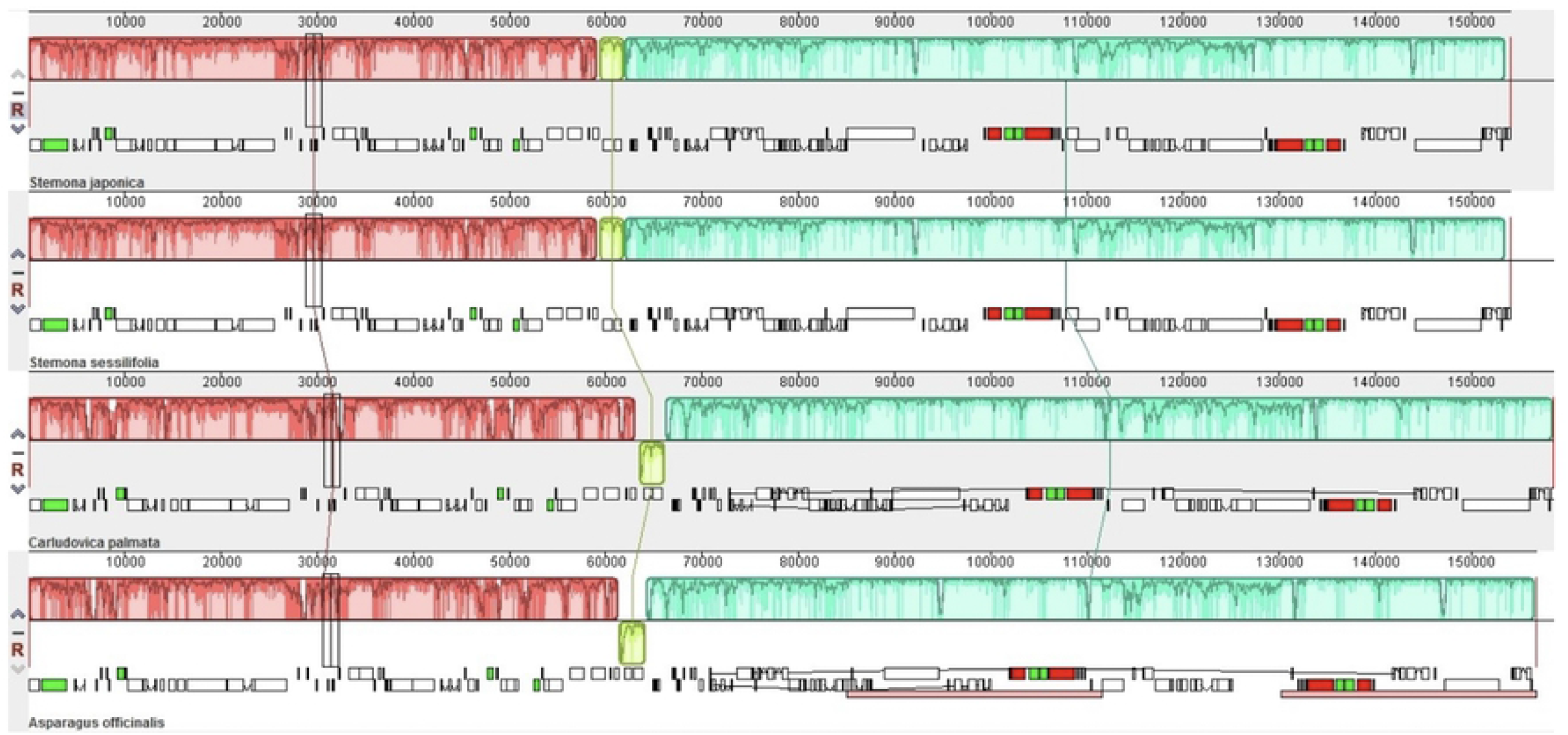
Comparison of three chloroplast genomes using MAUVE algorithm. Local collinear blocks were colored to indicate syntenic regions, and histograms within each block indicated the degree of sequence similarity.

### IR expansion and SSC contraction

IR contraction and expansion are common evolutionary events contributing to chloroplast genomes size variation [43]. Here, boundary comparison analysis was performed by which we attempt to identify IR contraction and expansion events (Fig 5). Compared to *A. officinalis* and *C. palmate,* the relatively larger IR regions indicated IR expansion events in *S. sessilifolia.* Simultaneously, the SSC region was shorter than *A. officinalis* and *C. palmate* by 465-737bp, suggesting the occurrence of SSC contraction events in *S. sessilifolia.* For *A. officinalis* and *C. palmate, the rpl22* gene is located at the LSC region with one copy. However, the IR regions of *S. sessilifolia* spanned to the intergenic spacers between *the rpl22* gene and *rps3* gene, resulting in two copies of the *rpl22* gene. Therefore, we can claim that the significant difference in *rpl22* coding regions between *S. sessilifolia* and its closely related species was attributed to IR expansion events. The IRb/SSC boundary extended into the ycf1 genes by 1146-1260bp, creating ycf1 pseudogene in *S. sessilifolia* and *C. palmate*. Considering premature stop codons were discovered in the ycf1 gene, only one ycf1 pseudogene was annotated in the SSC region in *A. officinalis*. The *ndhF* gene located at SSC regions in *A. officinalis* and *C. palmate,* and it ranges from 10-40bp away from the SSC/IRa boundary. However, in *S. sessilifolia,* the SSC region’s shortening leads to the *ndhF* gene extended into the IRa region by 186bp. The *ndhF* gene located at the SSC/IRa junction resulted in partial duplication of this gene at the corresponding region. An overlap of 186bp between the *ndhF* gene and *ycf1* pseudogene was also observed in *S. sessilifolia.* Summarily, compared to *A. officinalis* and *C. palmate,* significant boundary expansion and contraction events were observed in *S. sessilifolia* simultaneously.

**Figure 5.**
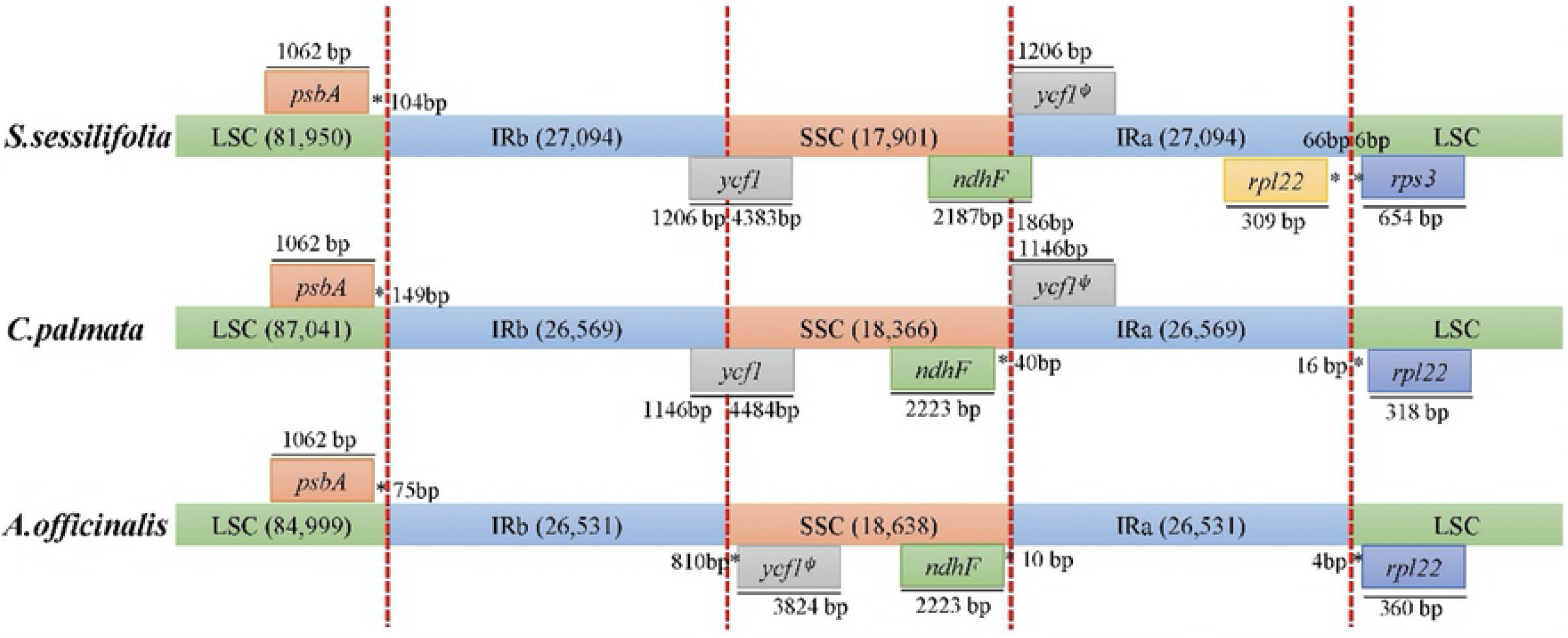
Comparison of IR, LSC, and SSC regions among *Stemona sessilifolia, Carludovica palmata,* and *Asparagus officinalis.* Numbers around the genes represented the gene lengths and the distances between the gene ends and boundary sites. Please note that the figure features were not to scale. ^ψ^ indicates pseudogene.

### Phylogenetic Analysis

The chloroplast genome has been successfully used to determine plant categories and reveal plant phylogenetic relationships [44, 45]. To determine the phylogenetic position of *S. sessilifolia,* we constructed a phylogenetic tree with species in Stemonaceae and its closely related families (Asparagoideae, Velloziaceae, Cyclanthaceae, Pandanaceae). A total of 13 chloroplast genomes were retrieved from the RefSeq database, and 58 protein sequences shared by these species were used to construct a phylogenetic tree with *Veratrum patulum*, and *Paris dunniana* served as an outgroup (Fig 6). As a result, species in Stemonaceae, Asparagoideae, and Velloziaceae formed a cluster, respectively. Besides, *S. sessilifolia* and *S. japonica* formed a cluster within Stemonaceae with a bootstrap value of 100%, indicating the sister relationship between these two species.

**Figure 6.**
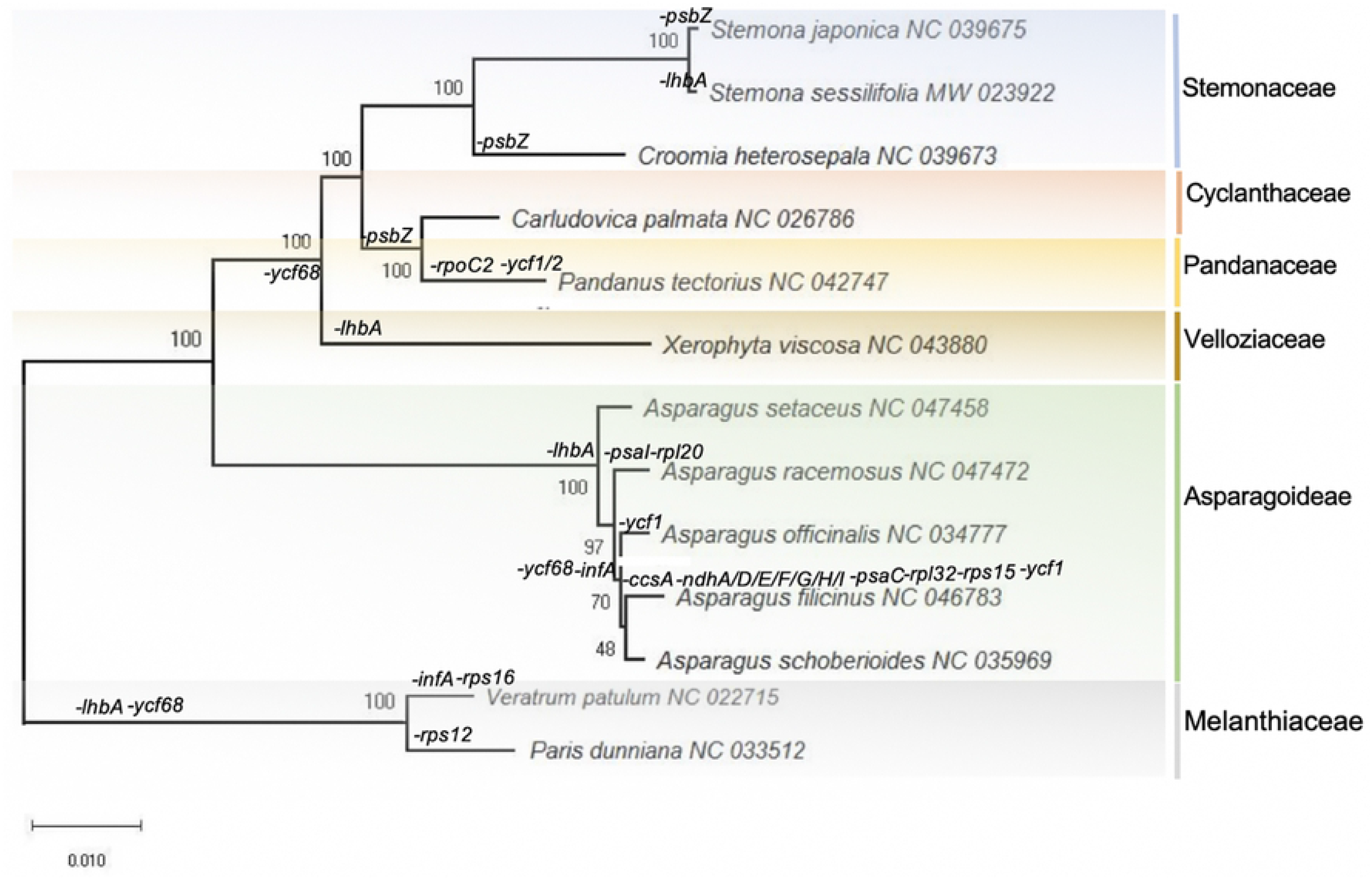
Molecular phylogenetic Analyses of Pandanales and its closely related orders. We constructed the tree with the sequences of 58 proteins presented in 116 species using the Maximum Likelihood method implemented in RAxML with *Nicotiana tabacum* and *Solanum Lycopersicum* served as an outgroup. The numbers associated with the nodes indicate bootstrap values tested with 1000 replicates. We marked the orders and families for each species besides the branches and the occurrence of gene loss events.

As showed in Fig 6, a series of gene loss events were observed throughout Stemonaceae and its closely related families (Asparagoideae, Velloziaceae, Cyclanthaceae, Pandanaceae). A total of 21 genes are lost in these species, including *ycf68* (11), *lhbA* (9), *infA* (4), *psbZ* (4), *ycf1* (3), *ccsA* (1), *ndhA* (1), *ndhD* (1), *ndhE* (1), *ndhF* (1), *ndhG* (1), *ndhH* (1), *ndhI* (1), *psaC* (1), *psaI* (1), *ycf2* (1), *rps16* (1), *rpl20* (1), *rpoC2* (1), *rps12* (1), and *rps15* (1), the number enclosed in parentheses represented the frequency of gene loss events. As expected, closely related species always tend to undertake the same gene loss events. A series of clusters formed by species undertaken the same gene deletion events further confirmed such a phenomenon. *C. palmata* and *P. tectorius* formed a cluster that lacked psbZ gene. The species from Pandanales (Steminaceae, Cyclanthaceae, Pandanaceae, and Velloziaceae) formed a cluster without ycf68 gene. The species from Asparagoideae formed a cluster without *lhbA* gene.

*Ycf68* gene has the highest frequency of gene deletion, and the second was *lhbA* gene. The following three were *the infA* gene, *psbZ*, and *ycf1* gene, respectively. The *ycf68* gene was only found in two species *(Asparagus racemosus, Asparagus setaceus),* and *the lhbA* gene was only found in four species (*C. palmata, C. heterosepala, P. tectorius* and *S. japonica).* The function of *the ycf68, lhbA,* and *ycf1* gene remained unknown. The occurrence of premature stop codons may account for these three genes’ rare existence in chloroplast genomes [38, 46, 47]. As one of the most active genes in the chloroplast genome, *the infA* gene plays an essential role in protein synthesis. The frequent absence of the *infA* gene may contribute to the transfer of this gene between cytoplasm and nucleus [38, 48]. The lack of the subunits of the photosystem II gene psbZ was frequently observed in Pandanales (Steminaceae, Cyclanthaceae, Pandanaceae). For each of the remaining 16 genes, only one gene loss event was observed, respectively. There was gene absence in each species’ chloroplast genome, indicating the variation in chloroplast genomes’ contents. However, for 16 out of 21 genes, the frequency of gene loss events was only one, suggesting the chloroplast genome is highly conserved on the scale of gene contents. Such a phenomenon is consistent with the highly conserved nature of the chloroplast genome and its feature of rich in variation.

## Discussion

In this study, we sequenced and analyzed the chloroplast genome of *Stemona sessilifolia* and performed multi-scale comparative genomics of *Stemona sessilifolia, Asparagus officinalis,* and *Carludovica palmate* (the major counterfeit of Baibu). We also characterized the major changes in the chloroplast genome of *Stemona sessilifolia* compared with those of Dioscoreales, Liliales, and Pandanales, including genome rearrangement, IR expansion, and SSC contraction, and investigate the occurrence of gene loss events in Dioscoreales, Liliales, Pandanales, and Asparagaceae.

Our results show that the genome of *Stemona sessilifolia* is very similar to that of *Stemona japonica previously reported.* In both chloroplast genomes of *S. sessilifolia* and *S*. *japonica,* the rps12 gene contained two introns. It is a trans-spliced gene with a 5’ end exon located in the LSC region, and the 3’ end exon and intron located in the IR regions [49]. Also, we detected a large inversion in both species. The SSC region was found to have a reverse orientation in S. japonica. The SSC region’s reverse direction has been interpreted as a major inversion existing within the species [50–52].

Interestingly, a 3-kb long inversion was detected in the chloroplast genome of *S. sessilifolia.* It might result from a genome rearrangement event. This unique inversion phenomenon led to significant differences in the ycf3-psbJ region between *Stemona sessilifolia* and its related species, which can be used as a candidate region to identify *Stemona sessilifolia* from counterfeits.

SSRs have been widely used as molecular markers in the studies of species identification, population genetics, and phylogenetic investigations based on their high-degree variations [53]. The SSR consisting of A/T is the *most abundant type in S. sessilifolia and S. japonica.* These SSRs loci were mainly located in intergenic regions and would help develop new phylogenetic markers for species identification and discrimination [49]. Only forward and palindrome repeats were found in *the S. sessilifolia* cp genome regarding the long repeat sequences. The biological implication of these repeats remains to be elucidated.

Also, there were significant differences in IR contraction and expansion between *Stemona sessilifolia* and other species. At the IRa/LSC border, the spacer from rpl22 coding regions to the border is longer in *Stemona sessilifolia* (309 bp) than that of *Stemona japonica* (65 bp). The IRb/SSC boundary extended into the ycf1 genes by only 18bp and created a ycf1 pseudogene in *Stemona japonica* [49]. However, that region is 1146-1260bp long in *Stemona sessilifolia.* The function of ycf1 genes is mostly unknown, but it evolves rapidly [54]. The larger contraction and expansion of the IR region in *Stemona sessilifolia* may lead to evolutionary differences between *Stemona sessilifolia and* its closely related species. This may need further verification.

*Stemona sessilifolia* and *Stemona japonica* are the authentic sources of Baibu, according to Pharmacopoeia of the People’s Republic of China (2015 edition). Phylogenetic analyses showed that they were placed close to each other with a bootstrap value of 100%. *Asparagus officinalis* and *Carludovica palmate* (the major counterfeit of Baibu) were on the other branches. When we investigated the gene loss events in the phylogenetic relationship context, we also see the cp genomes of *Stemona sessilifolia* and *Stemona japonica* have similar gene loss patterns. These findings support the pharmaceutical use of *Stemona sessilifolia* and *Stemona japonica as* genuine Baibu. Also, they suggest the urgent need for new molecular markers for the identification of genuine Baibu. This study will be of value in determining genome evolution and understanding phylogenetic relationships within Pandanales and other species closed to Pandanales.

## Conclusions

In summary, the complete plastome of *Stemona sessilifolia (Miq.) Miq.* was provided in the current study. We believe it will benefit as a reference for further complete chloroplast genome sequencing within the family. A multi-scale comparative genome analysis among *Stemona sessilifolia, Asparagus officinalis,* and *Carludovica palmate* (the major counterfeit of Baibu) was based on sequence data provided performed. Comparative Analysis of these three species revealed the existence of a unique inversion in the ycf3-psbJ regions. Interestingly, IR expansion and SSC contraction were observed simultaneously in *Stemona sessilifolia*, resulting in a rare boundary pattern. some highly variable regions were screened as potential DNA barcodes for identification of these three species, including *trnF-GAA_ndhJ, atpB_rbcL, rps15_ycf1, trnG-UCC_trnR-UCU, ndhF_rpl32.* Phylogenetic analyses showed that the two Stemona species were placed close to each other with a bootstrap value of 100%. Finally, we investigated the gene loss events in the context of the phylogenetic relationship. Closely related species always share similar gene loss patterns, consistent with those observed previously. This study will be of value in determining genome evolution and understanding phylogenetic relationships within Stemonaceae and families closed to Stemonaceae.

## Acknowledgments

This work was supported by the CAMS Innovation Fund for Medical Sciences (CIFMS) (2016-I2M-3-016, 2017-I2M-1-013) from the Chinese Academy of Medical Science, National Natural Science Foundation of China (31460237),Guangxi science and technology program (AD17292003), Guangxi R&D project of medical and health technology (S201529). The funders were not involved in the study design, data collection, and Analysis, decision to publish, or manuscript preparation.

## Author Contributions

CL and *WW* conceived the research; JTL and MJ carried out the bioinformatics studies and prepared the manuscript; HMC and YL collected samples of *Stemona sessilifolia,* extracted DNA for next-generation sequencing. All authors have read and approved the manuscript.

## Supporting information

**S1 File. The barcode sequences of *Stemona sessilifolia* available in GeneBank.**

**S2 File. The sequence of *Stemona sessilifolia, Carludovica palmata,* and *Asparagus officinalis.***

**S1 Table. List of chloroplast genomes used in this study.**

**S2 Table. The length of introns and exons for intron-containing genes.**

**S3 Table. K2p distances for intron regions among *Stemona sessilifolia, Carludovica palmata*, and *Asparagus officinalis*.**

**S4 Table. K2p distances for intergenic regions among *Stemona sessilifolia, Carludovica palmata*, and *Asparagus officinalis*.**

